# Fitting the Balding-Nichols model to forensic databases

**DOI:** 10.1101/009969

**Authors:** Rori V. Rohlfs, Vitor R. C. Aguiar, Kirk E. Lohmueller, Amanda M. Castro, Alessandro C. S. Ferreira, Vanessa C. O. Almeida, Iuri D. Louro, Rasmus Nielsen

## Abstract

Large forensic databases provide an opportunity to compare observed empirical rates of genotype matching with those expected under forensic genetic models. A number of researchers have taken advantage of this opportunity to validate some forensic genetic approaches, particularly to ensure that estimated rates of genotype matching between unrelated individuals are indeed slight overestimates of those observed. However, these studies have also revealed systematic error trends in genotype probability estimates. In this analysis, we investigate these error trends and show how the specific implementation of the Balding-Nichols model must be considered when applied to database-wide matching. Specifically, we show that in addition to accounting for increased allelic matching between individuals with recent shared ancestry, studies must account for relatively decreased allelic matching between individuals with more ancient shared ancestry.

## 1. Introduction

Forensic databases, rapidly increasing in size, invite powerful analyses of rates of coincidental genotype matching [1, 2, 3]. Such analyses have validated some basic assumptions in forensic genetics, particularly the reasonable over-estimation of genotype frequencies with existing methods. However, these studies also illustrate how database population genetic diversity differs from what is expected under the basic model of forensic genetics: the Balding-Nichols (BN) model.

The BN model simply and elegantly provides a framework for estimating probabilities of observed genotypes, taking into account population structure and variance in allele frequency estimates [4, 5]. The BN model can be interpreted as describing an ancestral population which has split into a number of internally randomly mating sub-populations which evolve independently over some time, resulting in a present-day total population made up of a number of cryptic sub-population groups. The sampling probabilities estimated under the BN model then incorporate the deviations from Hardy-Weinberg equilibrium expected due to population divergence.

The amount of excess allele-sharing in a sub-population group beyond what is expected based on the total population allele frequencies can be quantified in the BN model by the parameter *θ*. *θ* can be thought of as the probability that two alleles in a sub-population are identical by descent (IBD) to due to within sub-population shared ancestry. In a coalescent framework under simplifying assumptions, it represents the probability that two alleles sampled from within a sub-population coalesce before either mutates or migrates out of the sub-population [4].

In the BN model used in forensic applications, the probability of observing a particular genotype conditioning on having observed the same genotype is estimated using the *θ* correction to account for coincidental allelic sharing between two individuals due to excess shared ancestry within a sub-population. In most forensic calculations, there is an implicit assumption that the individuals in question are from the same sub-population [4]. Balding and Nichols convincingly argue that this assumption is appropriate, saying “the ‘same sub-population’ assumption is conservative, since the suspect’s profile will tend to be more common in his/her sub-population than in other groups” [4]. A number of studies have shown the importance and appropriateness of this assumption and the corresponding *θ* correction in genetic identification calculations [5, 6, 7, 8, 9, 10, 11, 12].

In database applications, typically all pairs of genotypes in a database will be compared to each other and their degree of matching assessed. Previous applications of the standard BN model to forensic databases [1, 2] have shown that the often-used *θ* correction of 0.01 usually adequately corrects for coincidental allele-sharing, raising estimated probabilities of matching genotypes above their observed levels, and therefore reducing false positive rates below their expectation (in statistical terms, making the test ‘conservative’). Yet, these analyses show an excess of non-similarity between observed pairs of individuals, as compared to the expectation [1, 2]. As we will show, this is likely due to the fact that the standard formulation of the BN model does not take decreased allele sharing between individuals from different sub-populations into account. When applying the BN model to describe the amount of genotypic matching observed in a database, it is not clear that the ‘same sub-population’ assumption is appropriate.

In this manuscript, we investigate how empirical genotype matching observations can be explained by reconsidering the implementation of the BN model. We show that by accounting for the case of two individuals deriving from different population groups, we significantly improve the ability to describe empirical matching rates in a database.

## 2. Methods

### 2.1. Allele sharing matrix

To quantify the degree of multi-locus genotype matching within a data set, consider the matrix *M* where each entry *M*_*m,p*_ is the number of profile pairs with *m* markers matching at both alleles and *p* markers matching at one allele [1, 2]. Tvedebrink *et al.* [2] described a recursive algorithm to compute the probability *π_m,p_* that two multi-locus genotypes completely match at *m* loci and partially match at *p* loci, constructing a probability matrix *π* analagous to *M*. This method uses the single-locus probabilities of individuals matching two, one, and zero alleles as *P*_1,0_, *P*_0,1_, and *P*_0,0_, respectively, following in the notation of Tvedebrink *et al.* [2]. Note the parallel notation to counts of matching and partially matching markers in *π_m,p_*. Weir [1] described how to compute *P*_1,0_, *P*_0,1_, and *P*_0,0_ at a locus by summing over the appropriate two-individual single locus genotype probabilities [1].

### 2.2. Single locus allelic sharing probabilities

#### 2.2.1. Individuals from the same sub-population group

Under the typical implementation of the BN model, where all individuals are assumed to be in the same sub-population group, the two-individual genotype probabilities are

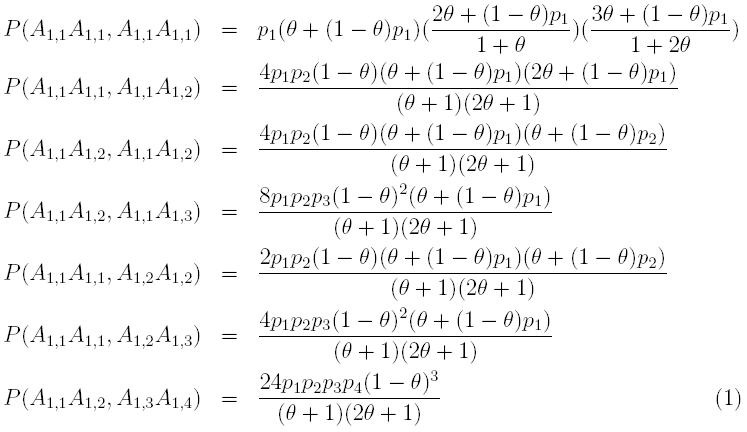

where *A*_1*,i*_ is an allele *i* drawn from the single sub-population 1, so, for example *P* (*A*_1*,i*_*A*_1*,i*_*, A*_1*,i*_*A*_1*,j*_) is the probability observing a homozygote and heterozygote sharing one allele, and *p*_*i*_ is the frequency of allele *i*.

#### 2.2.2. Individuals from same or different sub-population groups

Under the BN model, if two individuals are not in the same population group, the probability that their alleles coalesce more recently than a mutation or migration event is zero. In other words, there is no increased chance of allele-sharing due to shared ancestry for individuals in different population groups. In that case, the probability of observing their genotypes is computed as a function of the observed allele frequencies without the *θ* correction.

We can allow individuals to be from different sub-populations by introducing a parameter *d*, which describes the probability that a pair of individuals are from different sub-population groups. This way, we fully describe the BN model with some individuals from the same sub-population group and some from differing groups. Under a model with population differentiation, two-individual genotype probabilities are

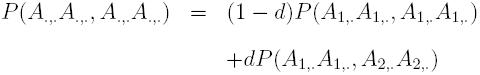

where subscript dots indicate any option such that *A*_.,._ is any allele drawn from any sub-population and *A*_1_,. is any allele drawn from sub-population 1. Genotype probabilities for individuals from the same population are the same as under the typical implementation of the BN model and for individuals from different sub-populations the genotype probabilities are

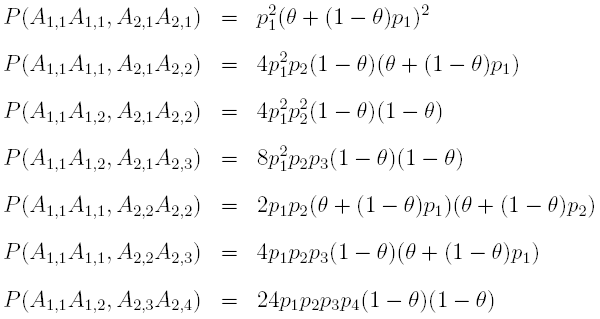

#### 2.2.3. Chromosomes from same or different sub-populations

In the previous formulation of joint genotype probabilities for two individuals, it is assumed that in each individual, both chromosomes derive from the same sub-population. In post-colonial societies, where few individuals can trace all their ancestry to their current location, this is not realistic. We describe an alternative model allowing alleles within individuals to be drawn from different, but correlated, sub-population groups. In this model there are *k* sub-populations of equal size and relation to each other. The correlation of sub-population draws within individuals is described by the parameter *a*. We can use this model to compute joint genotype probabilities, as shown in Supplemental Materials.

### 2.3. Likelihood framework

With match probabilities specified by the aforementioned models, we can calculate the expectation *π* of the match matrix *M* under varying assumptions regarding allele frequencies, and parameters of the models: *θ* for the typical implementation of the BN model without population differentiation, *θ* and *d* for the model with population differentiation, and *θ*, *a*, and *k* for the model allowing admixture between sub-populations (Table 1). By taking the entries *π* as categorical probabilities in a multinomial distribution, we can compute the sampling probability of an observed instance of *M*, an approach used effectively in other population genetic applications [13].

**Table 1:** The maximum log likelihoods and parameter estimates are listed for each model considered. For models which do not incorporate particular parameters, those estimates are listed as NA.

| model | log likelihood | $\hat{\theta}$ | $\hat{d}$ | $\hat{a}$ | $\hat{k}$ |
| --- | --- | --- | --- | --- | --- |
| typical implementation with $\theta = 0.01$ | -47246554388 | NA | NA | NA | NA |
| typical implementation with $\theta$ varying | -1259283 | $4.6e - 10$ | NA | NA | NA |
| population differentiation implementation | -37234 | $7.6e - 03$ | 0.966 | NA | NA |
| sub-population groups by chromosome | -37182 | $7.9e - 03$ | NA | 0.939 | 3538.5 |

Using the sampling probability of *M* as a likelihood function, we can estimate parameters of the model using maximum likelihood. Since the models described here are nested and fulfill standard regularity conditions, the asymptotic distributions of likelihood ratio test statistics (LRTs) are known to be chi square. Specifically, if we take the null hypothesis to be the typical implementation of the BN model with a fixed value of *θ* (say *H*_0_ : *θ* = 0.01), and the alternative to be the typical BN model implementation where *θ* varies (*H*_*a*_ : *θ* ≠ 0.01), the LRT is distributed as a chi square with one degree of freedom 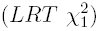. Similarly, to compare the typical implementation of the BN model with our implementation with population differentiation, we specify *H*_0_ : *θ* ≠ 0.01*, d* = 0 and *H*_*a*_ : *θ* ≠ 0.01*, d* ≠ 0, in which case the LRT 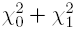. The model allowing chromosomes within individuals from different sub-populations reduces to the model with population differentiation under complete allelic correlation (*a* = 1). In this case, *d* is equivalent to (*k* − 1)*/k*. This enables tests where LRT 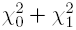 between the chromosomal model and the model with population differentiation

Additionally, we can obtain parameter estimates of 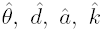 and *ĉ* While we do not advocate interpreting these estimates too strongly, as the underlying population models are very simple, we can compare them as a reference.

### 2.4. Database

We consider genotype data from 99,275 Brazilian individuals undergoing paternity testing during 2011-2013 in the Hermes Pardini Laboratory, Vespasiano, MG, Brazil. The individuals genotyped reside in all 26 Brazilian States and the Federal District (Brasilia). The genotypes were obtained using a combination of two Life Technologies kits and ABI 3730 Genetic analyzers (Life Technologies, CA, USA) for a total of 20 loci (the original 13 CODIS core loci and additionally D10S1248, D22S1045, D1S1656, D12S391, D2S441, D2S1338, D19S433, PentaD, and PentaE) [14].

While there are no known relatives in this dataset, unknown relatives, or multiple entries of the same individual are expected. As such, individuals with 17-20 loci matching and the same birth dates (when available) were removed as likely multiple entries or identical twins with some genotyping or clerical errors. When dates were not available or inconsistent apparently due to a typo, names were manually checked by the lab personnel and the most complete profile was kept, resulting in a data set with 96,400 individuals [14].

Since our analysis requires genotypes across the same number of loci for all individuals, we discarded all individuals with any missing data. In the remaining data set, extremely rare alleles observed exactly one time may be, in fact, genotyping errors. Profiles with these rare alleles were similarly eliminated. The final dataset considered in this analysis contained 90, 852 individuals.

## 3. Results

### 3.1. Observed database matching

We counted the number of zero, one, and two allele matches for each locus for each pair of individuals in the dataset to create the observed matrix *M*_*obs*_, as shown in Supplemental Table 1. For example, in our dataset, out of 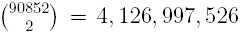 pairs of genotypes, 295, 948 pairs have exactly one loci matching at both alleles, two loci matching at one allele (partially matching), and 17 markers matching at neither allele (Supplemental Table 1).

### 3.2. Comparing data likelihood under different models

Previous investigators have used the conventional implementation of the BN model (without population differentiation) with *θ* fixed at 0.01 to describe matching in databases [1, 2]. Under this model, setting *θ* = 0.01, we calculated the log likelihood of the observed match matrix as -47, 246, 554, 388 (Table 1). We can graphically compare our observed and expected results in a dropping ball diagram [11, 15, 2] (Figure 1), or in a heatmap of the residuals (Figure 2a). The heatmaps in this manuscript show a color gradient along the log of the divergence of the observed and expected as 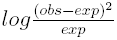. Through these visualizations, we see that as in previous analyses [1, 2], under the typical BN model implementation with *θ* set at 0.01, there is an excess of observed pairs of individuals who share few alleles, as compared to the expectation.

**Figure 1:**
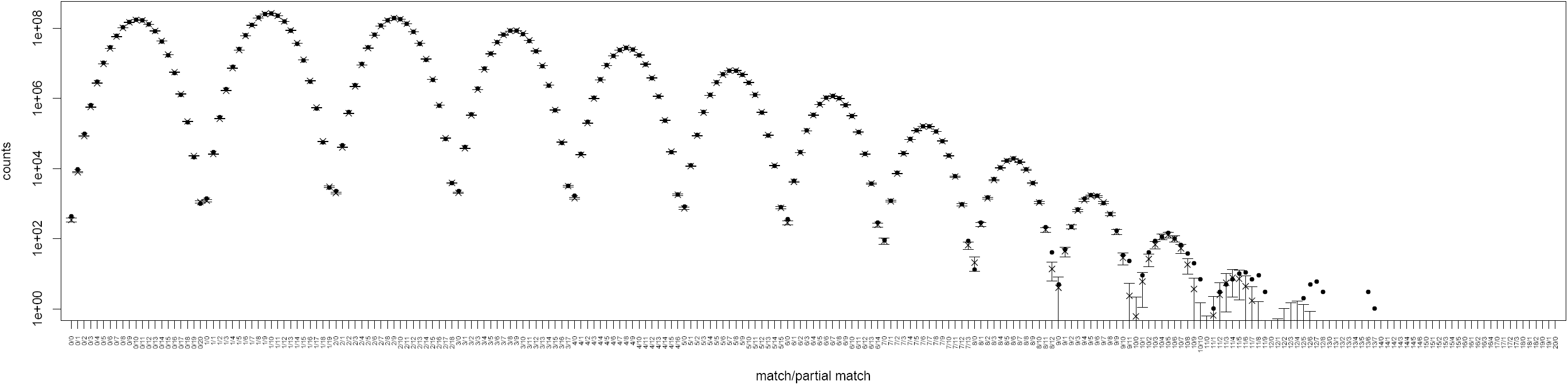
This dropping ball plot shows the observed (dot) and expected (x) numbers of pairs of individuals sharing *m* matching loci and *p* partially matching loci where *m/p* is indicated on the x-axis.

**Figure 2:**
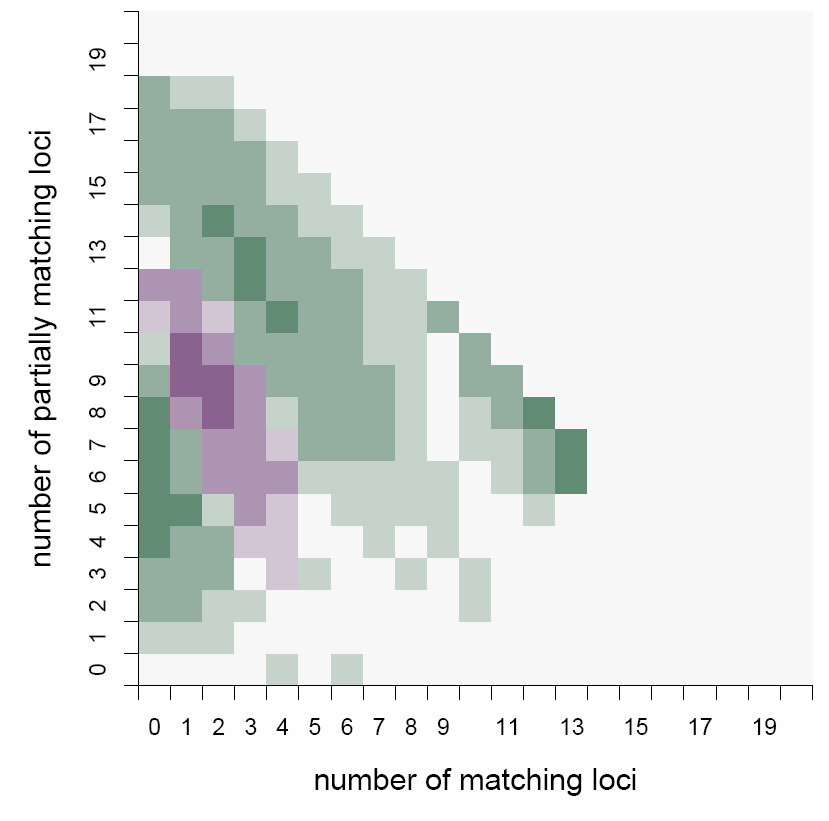
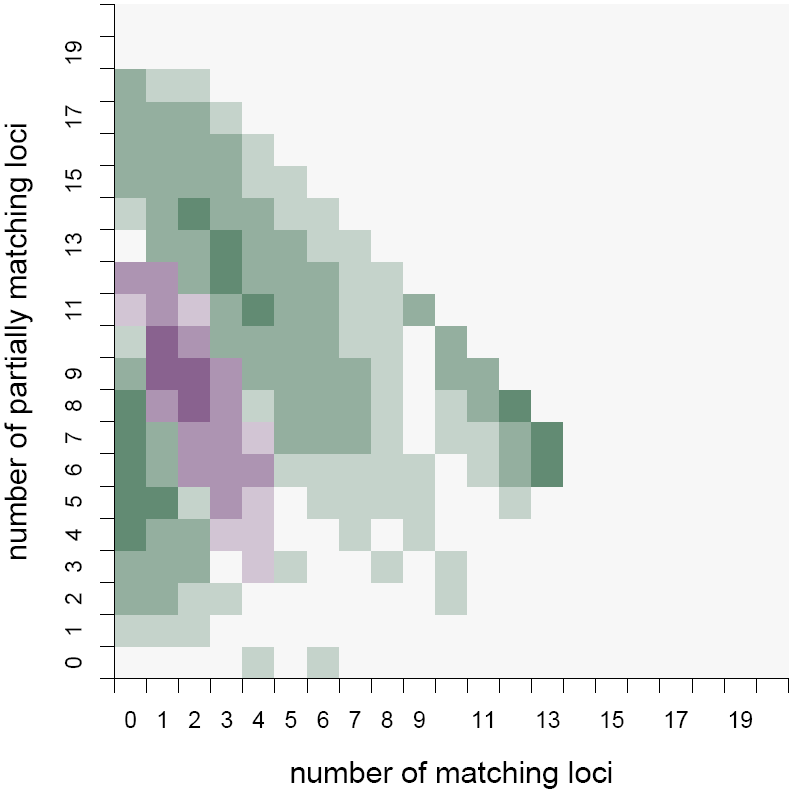
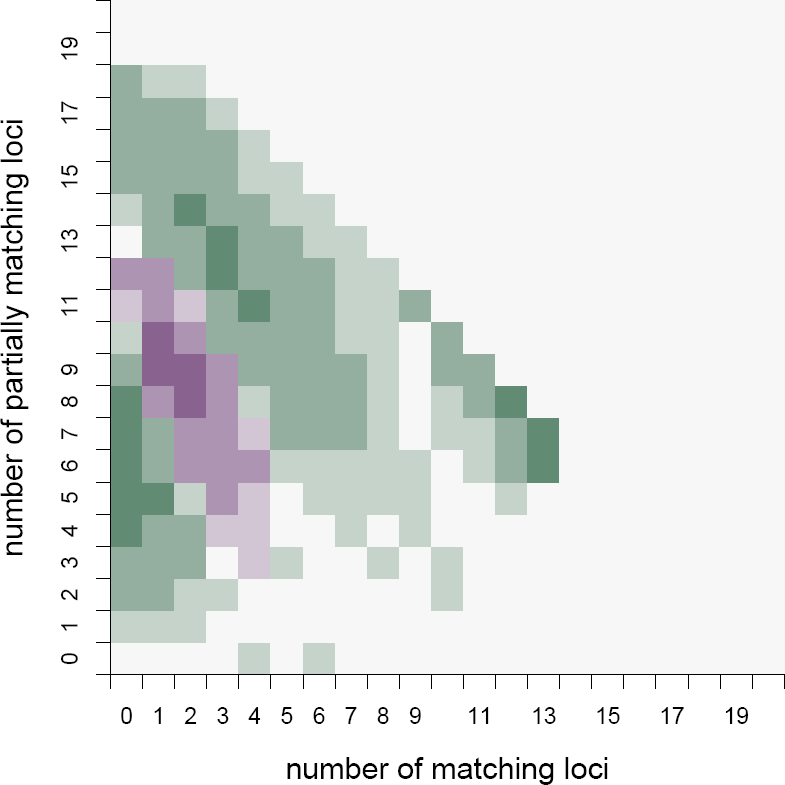
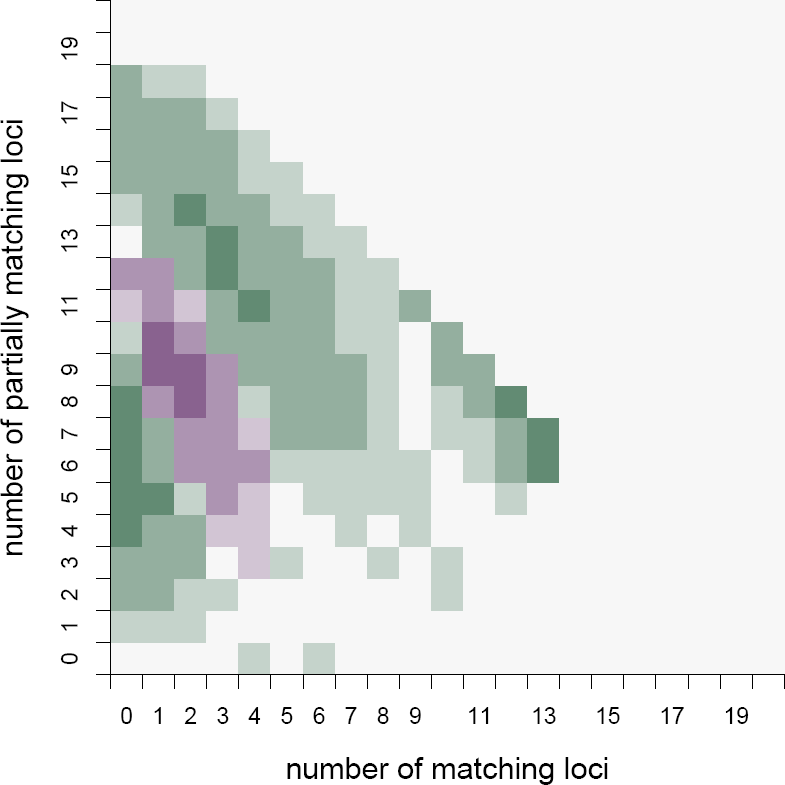
These heatmaps show the difference between the observed match matrix and that expected under, (a) the typical implementation of the BN model with *θ* = 0.01, (b) the typical implementation of the BN model where *θ* varies, (c) the full implementation of the BN model, and (d) the full implementation of the BN model allowing for admixture. Purple indicates a lack of observed pairs of individuals and green indicates an excess of observed pairs of individuals.

Using the maximum likelihood framework and optimizing over *θ*, we performed a similar analysis (Table 1, Supplemental Table 2). This model where *θ* may vary fits the observed data significantly better than with *θ* fixed at 0.01 (*LRT* = 94, 490, 590, 210). However, we still observed an excess of individuals sharing few alleles (Figure 2b). Further, under the maximum likelihood of this model, *θ* is estimated near zero as 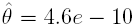, indicating that the *θ* correction as implemented here is insufficient to describe the data. This makes sense since the *θ* correction accounts for excess allelic sharing due to common ancestry within a sub-population.

We allow individuals in different sub-populations to share comparatively fewer alleles through common ancestry using the population differentiation model, where two random individuals derive from different sub-population groups with probability *d*. Again, we find the maximum likelihood value under this model (Table 1, Supplemental Table 3). The model fits the observed data significantly better (*LRT* = 2, 444, 098) and corrects for the previous excess of individuals sharing few alleles (Figure 2c). However, we still see consistent differences between the observed and expected allelic matching. Compared to the observed data, the population differentiation model predicts a more narrow range of allelic matching than what is observed.

In the population differentiation model, it is assumed that both alleles within an individual derive from the same sub-population. This assumption may not be valid in realistic cases of admixture, and can be relaxed using the a model where chromosomes are considered separately with some correlation. We fit such a model with *k* equally represented sub-populations and intra-individual allelic correlation to the observed data. This model, allowing chromosomes of different population origins within individuals, fits the data significantly better than the model without admixture (*LRT* = 148835) (Figure 2d). Still, we observe a wider range of allelic matching than is expected under these models.

## 4. Discussion and Conclusions

We have shown how a multinomial distribution on the expected match matrix can be used to calculate the sampling probability of an observed match matrix. Further, we have shown how this probability can be maximized with respect to some parameters to provide maximum likelihood estimates of these parameters.

Using this procedure, we found that estimating the value of *θ*, unsurprisingly, fits the data significantly better than a uniform value of 0.01. Further, we found that estimate to be near zero. This initially surprising estimate is explained by considering that the common implementation of the BN model in forensic genetics accounts for excess allele sharing due to recent ancestry, but not relatively less allele sharing for individuals with more distant ancestry. Under this implementation, every pair of individuals has increased allelic sharing due to recent ancestry. Since many pairs of individuals do not share recent ancestry, the maximum likelihood estimate of *θ* is driven to zero to explain the lack of consistent excess allele sharing.

We show and implement several parameterizations of the full BN model where individuals may or may not have excess allele sharing (equivalently, may or may not derive from the same population group). This full BN model fit the observed match matrix significantly better than the typical BN model implementation. Under the full BN model, 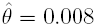, which is closer to the typical value used in practice of *θ* = 0.01.

Even under the full BN model implementation, we predict a more narrow range of locus-matching than observed. In the BN model, all sub-populations have equal excess allele sharing internally and are equally unrelated to each other. While this model provides a simple and reasonable over-estimation of coincidental genotype match rates, essential to forensic case work, it is clearly a simplification of complex human population structures, where some individuals are vastly more related than others. A more sophisticated model allowing varying degrees of allele sharing between individuals would likely better fit our observation of a broad range of allelic-matching. However, such a model would begin to accumulate parameters, making use in forensic case work impractical compared to the adequate typical BN model implementation.

Additionally, the full BN model does not explain a small observed excess of people matching at many loci. For example, there are three pairs of individuals who match both alleles at 13 loci and one allele at six loci, whereas under the full BN model, 5.0*e* − 13 are expected. There are several possible explanations for these individuals. They may be genetic relatives who share a large number of alleles IBD. They could share even more alleles than expected if allele frequencies are mis-specified because they derive from a population group divergent from the whole sample [16]. It is also possible that the same individual was entered a number of times, with genotyping or clerical errors resulting in differing alleles.

Other authors have considered the presence of genetic relatives within a database when calculating genotype match probabilities with additional parameters for the probabilities that a pair of individuals hold particular genetic relationships [1, 17, 2]. This way, the total probability of genotype matching takes into account the possibility of genetic relationships. However, since the loci are still treated independently, the small probability of a genetic relationship is factored in at each locus separately, rather than considering how genetic relatives share alleles across loci. As a result, unless there are extensive genetic relatives in a dataset, this does not dramatically affect the expected allelic matching.

We have shown how the correct full implementation of the BN model is crucial to understanding database-wide allelic matching. While this is essential for database applications, it does not affect forensic case work where the typical BN model implementation is adequate to reasonably overestimate the probability of coincidental genotype matching.

## Acknowledgements

We are immensely grateful to the individuals whose DNA samples were used in this study, without which none of this work would be possible. This work was supported in part by National Institutes of Health grant ???, National Science Foundation award 1103767, and a CAPES-Brazil scholarship (Programa Demanda Social and PhD Student Exchange Program BEX 8425/11-6) ??MORE??. The funders had no role in study design, data collection and analysis, decision to publish, or preparation of the manuscript. This study was approved by the Ethics Committee of the Federal University of Espirito Santo (Brazil) Health Sciences Department (No. 448327).

